# Exploiting rodent cell blocks for intrinsic resistance to HIV-1 gene expression in human T cells

**DOI:** 10.1101/2023.04.08.536105

**Authors:** Ryan T. Behrens, Jyothi Krishnaswamy Rajashekar, James W. Bruce, Edward L. Evans, Amelia M. Hansen, Natalia Salazar-Quiroz, Lacy M. Simons, Paul Ahlquist, Judd F. Hultquist, Priti Kumar, Nathan M. Sherer

## Abstract

HIV-1 virion production is inefficient in cells derived from mice and other rodents reflecting cell-intrinsic defects to interactions between the HIV-1 auxiliary proteins Tat and Rev and host dependency factors CCNT1 (Cyclin T1) and XPO1 (Exportin-1, also known as CRM1), respectively. In human cells, Tat binds CCNT1 to enhance viral RNA transcription and Rev recruits XPO1 to mediate the nuclear export of intron-containing viral RNA. In mouse cells, Tat’s interactions with CCNT1 are inefficient, mapped to a single species-specific residue Y261 instead of C261 in human. Rev interacts poorly with murine XPO1, mapped to a trio of amino acids T411/V412/S414 instead of P411/M412/F414 in humans. To determine if these discrete species-specific regions of otherwise conserved housekeeping proteins represent viable targets for inhibiting Tat and Rev function in humans, herein we recoded (“mousified”) each in human CD4+ T cells using precision CRISPR/Cas9-facilitated gene editing. Both edits yielded cells refractory to Rev or Tat activity, respectively, with isolated, isogenic CCNT1.C261Y cell lines remarkable in their capacity to exhibit near total inactivation of viral gene expression for all X4 and R5-tropic HIV-1 strains tested, and even the more distantly related lentiviruses including HIV-2 and SIV_agm_. These studies validate minor and naturally-occurring, species-specific differences in otherwise conserved human host factors as compelling targets for achieving broad-acting cell-intrinsic resistance to HIV’s post-integration phases.

**Importance:** Unlike humans, mice are unable to support HIV-1 infection. This is due, in part, to a constellation of defined minor, species-specific differences in conserved host proteins needed for viral gene expression. Here, we used precision CRISPR/Cas9 editing to engineer “mousified” versions of two of these proteins, CCNT1 and XPO1, in human T cells. CCNT1 and XPO1 are essential for efficient HIV-1 transcription and viral RNA transport, respectively, making them intriguing targets for gene-based inactivation of virus replication. Targeting either gene yielded antiviral phenotypes, with isogenic CCNT1-modified cell lines confirmed to exhibit potent, durable, and broad-spectrum resistance to HIV-1 and other pathogenic lentiviruses, and with no discernible impact on host cells. These results provide proof of concept for targeting CCNT1 (and potentially XPO1) in the context of one or more functional HIV-1 cure strategies.

## Introduction

Cell-intrinsic barriers to human immunodeficiency type 1 (HIV-1) replication can confer life-long resistance to HIV-associated diseases in humans. A hallmark example is a naturally occurring 32-bp deletion in the *CCR5* gene (*CCR5Δ32*) that reduces cell surface levels of CCR5, the surface receptor for transmitted “R5-tropic” forms of HIV-1 (1–3). Allogeneic bone marrow transplant using cells homozygous for *CCR5Δ32* have contributed to the only known instances of HIV/AIDS functional cure (4–7). These cases, known as the Berlin, London, and New York patients demonstrated that drug-free remission or cure is possible through targeted inactivation of CCR5. However, the potential drawbacks to gene-targeted CCR5-based cures include the potential for viral evolution of altered receptor tropism (8) (*e.g.,* outgrowth of CXCR4-tropic HIV-1) and the Δ32 allele being linked to increased risk of symptomatic flavivirus infections (9–11). These issues underscore the need to consider additional gene targets in the context of cell-based functional cures.

In this context, studies of HIV-1 replication in non-human mammals have revealed a trove of species-specific gene barriers to infection (12,13). For example, rhesus macaques resist HIV-1 due to the activities of endogenous APOBEC3G and TRIM5 proteins that disrupt intact viral genome delivery to targeted cells (14,15). Other forms of HIV-1 resistance can be attributed to species-specific differences to so-called host-dependency factors (HDFs); cellular proteins hijacked by the viral parasite to carry out specific life cycle stages. For example, mice and other rodents lack HIV-compatible versions of both CCR5 and the primary attachment receptor CD4, thereby resisting viral entry. Exogenous expression of human CD4 and CCR5 was shown to alleviate viral entry blocks in some rodent cell lines (16–28). However, these changes were insufficient to restore viral replication due to additional rodent cell blocks affecting HIV-1’s post-integration stages (29–37,32,38–50).

Of the post-integration rodent cell blocks, those affecting the essential HIV-1 accessory factors Tat and Rev are best characterized. In human cells, HIV-1 Tat drives viral transcription by recruiting the positive transcription elongation factor b (p-TEFb) complex, consisting of a heterodimer of the CDK9 kinase and its regulatory partner CCNT1, to the *trans*-activation response element (TAR) structure present in the 5’ portion of initiated viral transcripts (51). CDK9 phosphorylates the C-terminal domain of RNA polymerase II to promote processivity and synthesis of full-length viral RNAs (51). Human CCNT1 encodes a cysteine at residue 261 (C261) thought to coordinate a zinc ion that stabilizes the functional Tat/p-TEFb/TAR complex (34–36). The murine ortholog of CCNT1 encodes a tyrosine at the equivalent position (Y261), and is associated with poor Tat-dependent transcription (34–36).

By contrast, in human cells HIV-1 Rev enables the nuclear export of unspliced and partially-spliced viral RNAs that would otherwise remain retained in the nucleus (51). In the nucleus, Rev binds to and multimerizes on the Rev response element (RRE) structure present on unspliced and partially-spliced transcripts, and engages the XPO1 (Exportin-1, also called CRM1) nuclear export machinery (51). Human XPO1 features a species-specific surface element defined by a proline, methionine, and phenylalanine at residues 411, 412, and 414, respectively (P411, M412, and F414 in humans and T411, V412, and S414 in mice) that we and others have shown is needed for efficient Rev activity (48–50); likely important for the formation of a functional Rev/RRE/XPO1 export complexes consisting of two or more XPO1 export receptors (52,53). An alternative but non-mutually exclusive model suggests that Rev/RRE complexes bind with greater affinity to human XPO1 compared to the murine counterpart (54).

Based on this knowledge, in this study we asked if these discrete, naturally occurring, rodent-specific HDF blocks to HIV-1 Tat and Rev function could be engineered into human CD4+ Jurkat T cells, taking advantage of a recently developed precision CRISPR/Cas9 genome editing strategy. We provide strong evidence that both human *CCNT1* and *XPO1* can be “mousified”, *i.e.,* modified to engender the anticipated block, and as proof-of-concept in the context of an antiviral target, isolated isogenic *CCNT1.C261Y* cell lines that were effectively resistant to HIV-1 Tat activity and abolished viral gene expression despite the C261Y edit having no discernible effects on CCNT1-regulated host cell functions. Further, we demonstrate that CCNT1-C261Y cells are highly likely to exhibit cell-intrinsic resistance to all known HIV-1 strains and subtypes, inducing a state of forced viral latency consistent with a “block-and-lock” provirus inactivation scenario. Collectively, these results highlight the potential value of rodent-specific domains of *CCNT1* and *XPO1* as targets for anti-HIV intervention.

## Results

### A dual reporter strategy to screen for cell-intrinsic blocks to early (Tat-driven) and late stage (Rev-driven) HIV-1 gene expression

So that we could screen for cell-intrinsic blocks to Tat- and Rev-dependent HIV-1 gene expression, we first devised a flow cytometric assay to capture fluorescence profiles of cell populations infected with a two-color HIV-1 reporter virus (E-R-/2FP, Fig 1A); E-R-/2FP encodes *mCherry* from the *nef* locus to monitor early, Tat-dependent but Rev-independent gene expression and *mVenus* as a Gag fusion (MA-mVenus-CA) reporting on late, both Tat- and Rev-dependent gene expression. Inactivating mutations in *env* and *vpr* genes (E-/R-) limited virus replication to a single round and reduced cytopathic effects (Fig 1B).

**Fig 1.**
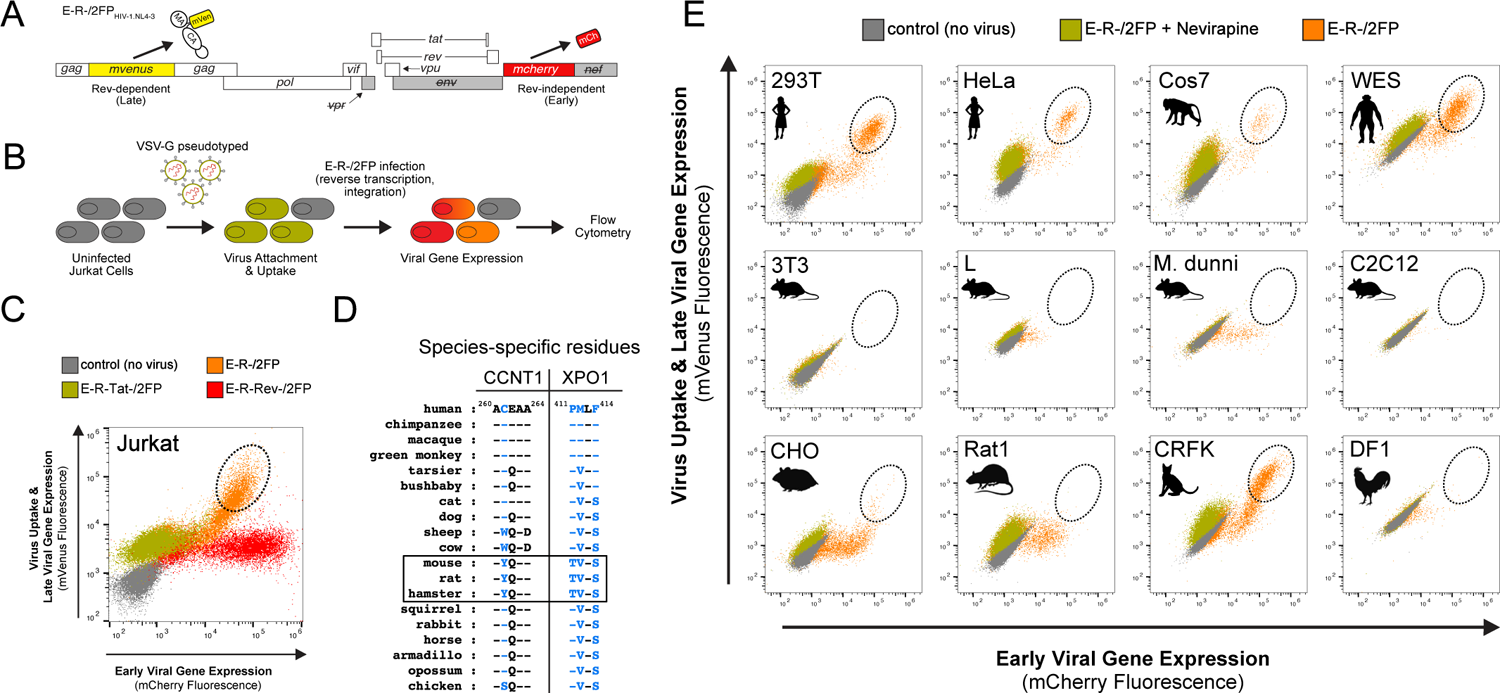
Development of a dual reporter assay to screen for blocks to early (Tat-driven) and late stage (Rev-driven) HIV-1 gene expression. (A) Viral genome diagram of the two-color, HIV-1.NL4-3-derived viral reporter construct, E-R-/2FP. Early, Rev-independent viral gene expression is reported by mCherry, encoded within the *nef* locus. Late, Rev-dependent viral gene expression is reported by mVenus, encoded in *gag* as an in-frame insertion between the matrix (MA) and capsid (CA) protein coding regions. Grey rectangles denote inactivated *env*, *vpr*, and *nef* genes to limit virus replication to a single cycle. (B) E-R-/2FP reporter assay schematic. VSV-G pseudotyped virus particles containing E-R-/2FP genomes are delivered to target cells. Prior to infection, target cells (grey) can become mVenus-positive due to virion-associated MA-mVenus during virus particle uptake (gold). Following proviral insertion, early (red) and late (orange) viral gene expression is reported by *de novo* mCherry and mVenus expression, respectively. (C) Fluorescence profiles of Jurkat E6-1 cells transduced with wild-type, Tat- or Rev-deficient E-R-/2FP, or untransduced control cells. Cells exhibiting robust early and late viral gene expression are indicated (dashed circle). Dot plot is an overlay of the four indicated cell populations. (D) Species-specific CCNT1 or XPO1 residues that govern HIV-1 Tat and Rev activity, respectively (blue). CCNT1 residue 261 governs Tat activity. XPO1 residues 411, 412, and 414 governs Rev activity. Rodents predicted from species-specific residue identity to exhibit poor Tat and Rev activities are indicated (boxed). (E) E-R-/2FP screen of various mammalian cell lines. Dashed circles highlight cells exhibiting robust early and late viral gene expression. Nevirapine treatment to block E-R-/2FP infection was to facilitate the identification of productively-infected cell populations.

To validate our reporter, we infected Jurkat T cells with our E-R-/2FP virus or control versions engineered to carry inactivating mutations in *tat* (E-R-Tat-/2FP) or *rev* (E-R-Rev-/2FP) (Fig 1C). Compared to mock-infected cells (grey dots), E-R-/2FP-infected cells (orange dots) yielded a robust population of bright double-positive cells (orange dots in dashed oval), consistent with efficient early and late viral gene expression phases. E-R-Rev-/2FP-infected cells (red dots) were mCherry-bright but mVenus-dim, consistent with Tat function but lacking Rev. As expected, E-R-Tat-/2FP-infected cells (gold dots) lacked both mCherry and mVenus fluorescence but like the E-R-Rev/2FP cells were moderately mVenus-positive due to cellular uptake of the Gag-mVenus fluorescent virus particles. In sum, E-R-/2FP control variants confirmed our dual reporter as capable of quantifying relative levels of Tat and Rev activity, with single cell resolution.

For further validation, we next tested our E-R-/2FP assay using a panel of informative cell lines predicted to encode permissive or non-permissive species-specific determinants encoded by *CCNT1* and *XPO1*, respectively (Fig 1D, shown in blue). Expectedly, primate cell lines including 293T, HeLa, Cos7, and WES exhibited fluorescence profiles similar to infected Jurkat T cells, confirming robust Tat and Rev activity (Fig 1E). In contrast, mouse cell lines including 3T3, L, and C2C12, exhibited deficiencies to Tat- and Rev-dependent gene expression, yielding few to no late stage, bright double-positive cells. Interestingly, M. dunni, Rat1 and CHO cells were largely deficient in late gene expression but exhibited a moderate degree of early, Rev-independent gene expression, illustrating that the transcription block was not absolute in these cell types despite their encoding CCNT1 Y261. Infected DF1 chicken cells, that encode a serine at the equivalent position of C261 were also only modestly permissive for early viral gene expression. CRFK cat cells were interesting in that, despite encoding a XPO1 patch region containing rodent-like V412 and S414 residues, these cells exhibited strong Rev activity, potentially highlighting the importance of XPO1 residue P411 or other yet to be mapped determinants. For all lines, treatment the reverse transcriptase inhibitor nevirapine (gold dots) served as a control to demarcate productively infected populations (orange dots) from uninfected cells (grey dots) (Fig 1E). Taken together, we concluded that our E-R-/2FP assay faithfully detected cell-intrinsic differences to Tat and Rev activities and would therefore be useful to screen modified human T cell populations for equivalent defects.

### CRISPR/Cas9 knock-in strategy to abrogate Tat or Rev function in human T cells

Next, we designed CRISPR/Cas9 gene knock-in schemes to determine if we could engineer rodent-informed Tat (CCNT1)- or Rev (XPO1)-dependent HIV-1 gene expression blocks into human CD4+ T cells, using the Jurkat E6-1 cell line as a tractable model system. Gene knock-in modifications were performed by co-delivering single-stranded oligodeoxynucleotide (ssODN) donor templates along with recombinant Cas9 in complex with guide RNAs bearing complementary sequences to targeted loci. For human *CCNT1*, the endogenous ‘TGC’ codon for cysteine at residue position 261 was replaced with ‘TAC’ in the ssODN, directing a substitution for the mouse-encoded tyrosine (Fig 2A, C261Y, modifications in red). Codon mutations were similarly made for combined proline, methionine, and phenylalanine residues at positions 411, 412, and 414 in human *XPO1*, to encode the mouse-encoded threonine, valine, and serine, respectively (Fig 2B, P411T, M412V, and F414S, modifications in red; collectively termed, “mPatch”). Silent mutations in ssODNs encoding novel restriction enzyme sites (BsiWI for *CCNT1* and PvuII for *XPO1*) allowed for knock-in detection using restriction enzyme digest (Figs 2A and 2B, modifications in blue). Protospacer adjacent motif (PAM) CRISPR targeting sequences were also mutated to prevent Cas9-mediated cleavage of introduced templates (Figs 2A and 2B, modifications in green).

**Fig 2.**
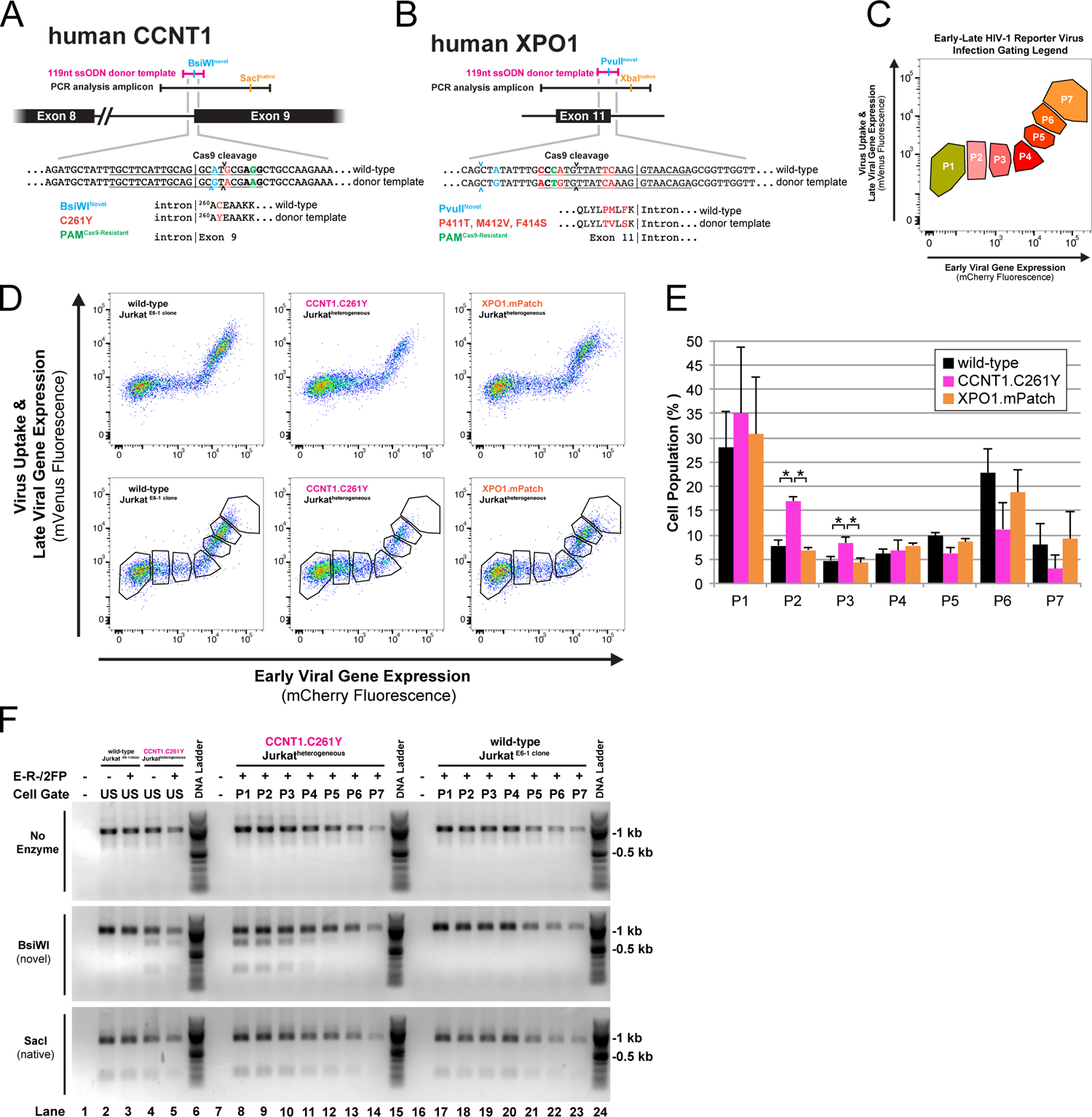
CRISPR/Cas9 knock-in strategy to abrogate Tat or Rev function in human T cells. (A) Design schematic for *CCNT1* gene knock-in editing. Approximate sequence lengths of the donor template (pink) and PCR amplicon (black) used to analyze the genomic locus. DNA sequences relevant to the edit design are shown; The PAM (bold) and guide RNA target (underlined) specify the Cas9 cleavage site (arrows). Mutations engineered into the donor template are color-coded according to their intended use. CCNT1 protein coding sequences are shown with substitutions color-coded to match the relevant DNA mutation. (B) Design schematic for *XPO1* gene knock-in editing labeled as in Fig 2A. (C) FACS gating legend for sorted populations P1-P7. Gates P1-P4 contain cells exhibiting early, Tat-dependent viral gene expression. Gates P5-P7 contain cells exhibiting late, Tat-/Rev-dependent viral gene expression. (D) Fluorescence profiles of *CCNT1- or XPO1-*edited or control cells infected with E-R-/2FP with (bottom row) or without (top row) P1-P7 gate overlays. P411T, M412V, and F414S substitutions collectively termed “mPatch”. (E) Percentage of cells inoculated with E-R-/2FP within gates P1-P7 for the *CCNT1*-(pink), *XPO1*- (orange), and control-edited (black) cell populations. Error bars represent the standard deviation from the mean for three independent experiments. A two-tailed Student’s *t*-test with Welch’s correction was used to compare the indicated populations (\**p* ≤ 0.05). (F) Sensitivity of PCR amplicons from E-R-/2FP-inoculated, *CCNT1*-edited or control cells before and after FACS. PCR amplicons (top row) were digested with BsiWI (middle row) or SacI (bottom row) to screen for amplicons containing gene knock-in for each sorted population (P1-P7, lanes 8-14 and 17-23). Unsorted E-R-/2FP-inoculated or no infection control cell populations (US, Lanes 1-5) were analyzed to demonstrate input amplicon sensitivities to the indicated restriction enzymes.

After editing, Jurkat cell populations were screened using the E-R-/2FP assay described for Fig 1. We hypothesized that, if refractory, *CCNT1*- or *XPO1*-modified cells would exhibit profiles similar to those observed for rodent cell lines (see Fig 2). To better resolve changes, we modified the assay to perform fluorescence-associated cell sorting (FACS) using seven cell gates (Fig 2C, bottom row; detailed in Fig 2D) allowing us to separate cell populations to represent the progression of viral gene expression beginning with mCherry-positive, Tat-dependent but Rev-independent stages (P1-P4) and ending with dual positive, late stage Rev-dependent stages (P5-P7). Comparing *CCNT1*-targeted cells to both wild-type and *XPO1*-edited cells, we observed marked increases to the number of cells in P2 and P3 gates, consistent with blocks to Tat function in a significant subset of *CCNT1*-edited cells (Fig 2E). While the number of cells in any gate of the *XPO1*-targeted cell population was not significantly different from wild-type, cells in gates P4-P7 were consistently brighter in the mCherry channel, consistent with Rev inactivation (Fig S1A, fluorescence ratio quantification in Fig S1B).

Because spontaneous insertion or deletion events can manifest off-target effects, we also analyzed each sorted pool to confirm allele knock-in frequency using restriction digest (P1-P7). Because the *CCNT1* knock-in was predicted manifest a Tat block, we hypothesized that alleles bearing the *CCNT1* knock-in signature would be elevated in cells within gates P1-P4 compared to those in the P5-P7 gates, confirmed by BsiWI sensitivity in the heterogeneous amplicon pools (Fig 2F). Analysis of *XPO1* knock-in by PvuII digestion sensitivity confirmed modifications, although for reasons unknown we observed no allele enrichment in any gated population (Fig S1B). Overall, these results indicated that mouse-encoded HIV-1 blocks could be successfully engineered into human T cells, with the single residue CCNT1.C261Y modification being relatively efficient.

### *CCNT1.C261Y* Jurkat T cells suppress HIV-1 Tat activity

Data indicating strong and stable blocks to Tat function in *CCNT1*-edited Jurkat cells prompted us to isolate two *CCNT1.C261Y* cell lines, named C1 and C2, encoding *CCNT1* loci confirmed to be entirely sensitive to BsiWI digestion (Fig 3A) and seamlessly edited, confirmed by the absence of unintended mutations in the targeted locus (Fig 3B and Fig 3C). CCNT1.C261Y protein levels were equivalent to non-edited CCNT1 in the parental cell line (Fig 3D) and we observed no effects on cell proliferation (Fig 3E). The E-R-/2FP reporter assay confirmed that the modified cells exhibited a fluorescence profile consistent with loss of Tat function (Fig 3F, compare panels i. to iv. & vii; compare all to Fig 1C) and, importantly, significantly rescued by restoring exogenous expression of human CCNT1 (using a retroviral vector) (Fig 3F and Fig 3G).

**Fig 3.**
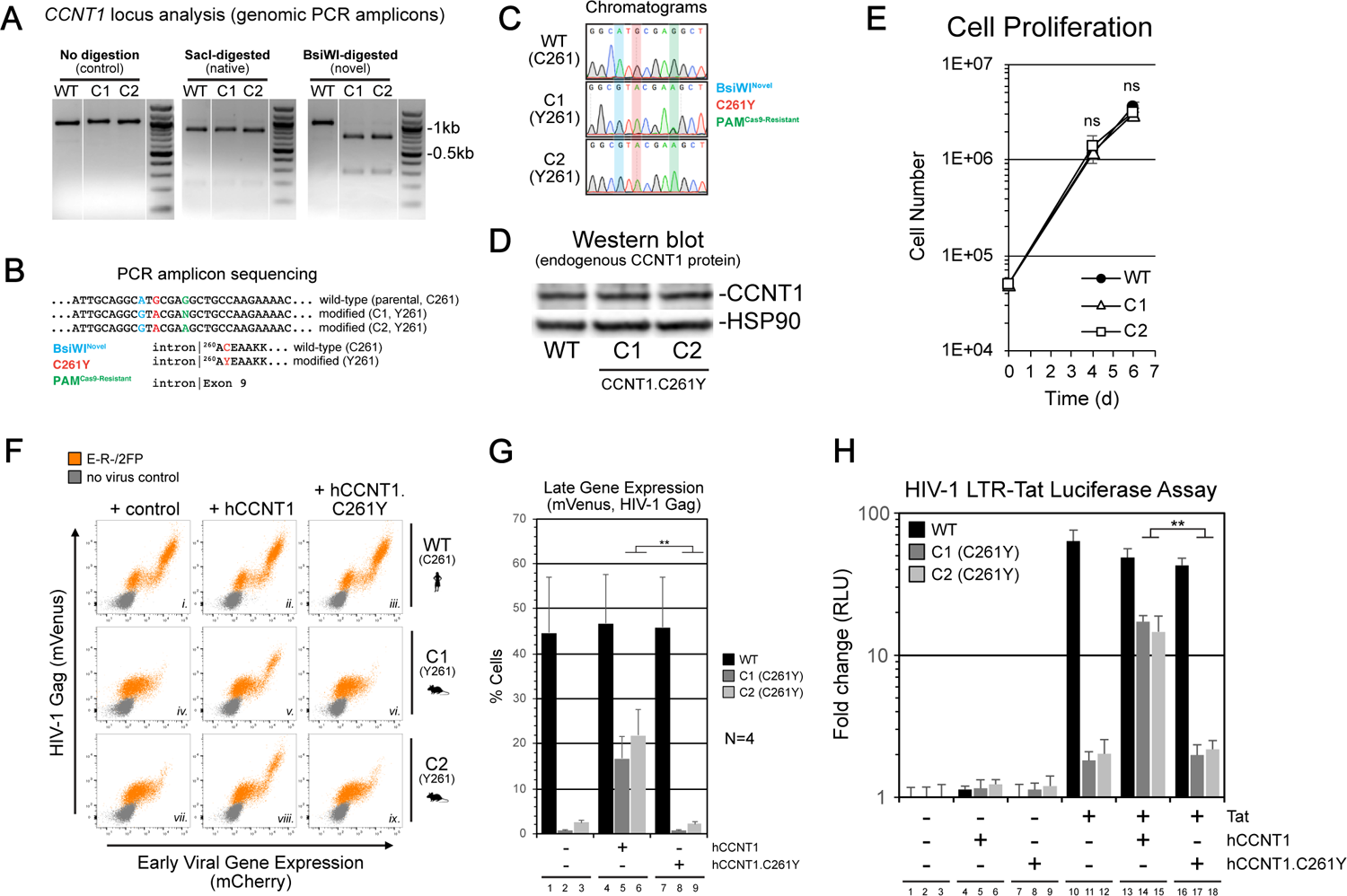
Homozygous CCNT1.C261Y Jurkat T cells potently suppress HIV-1 Tat activity. (A) Analysis of the *CCNT1* locus by PCR amplicon restriction digest for CCNT1.C261Y clones, C1 and C2, compared to WT parental cells. (B) Sequence analysis summary for CCNT1.C261Y cells. (C) Sequence read chromatograms for CCNT1.C261Y cells. Read peaks of loci targeted for mutation are color-coded by intended purpose. (D) Western blot analysis of CCNT1 protein levels in CCNT1.C261Y cells. (E) Cell proliferation analysis of CCNT1.C261Y cells. A two-tailed Student’s *t*-test with Welch’s correction was used to compare C1 or C2 to wild-type parental cells at the specified time point (ns, not significant). (F) Exogenous hCCNT1, but not hCCNT1.C261Y, restores the E-R-/2FP gene expression and the wild-type fluorescence profile in CCNT1.C261Y clones, C1 and C2. Vectors encoding wild-type hCCNT1 (middle column) or hCCNT1.C261Y (right column) or control (left column) were delivered to the parental Jurkat cell line (top row, panels i.-iii.) or CCNT1.C261Y clones C1 (middle row, panels *iv.-vi.*) and C2 (bottom row, panels *vii.-ix.*) then infected with E-R-/2FP virus (orange) and analyzed by flow cytometry. (G) Quantitative summary of late gene expression presented in Fig 3F. A two-tailed Student’s *t*-test with Welch’s correction was used to compare CCNT1.C261Y cells expressing exogenous hCCNT1 to those expressing hCCNT1.C261Y. (H) Exogenous hCCNT1, but not hCCNT1.C261Y, restores Tat activity in an infection-independent Tat/LTR-luciferase reporter assay. Plasmids encoding luciferase under control of the HIV-1 LTR and HIV-1 Tat were delivered in combination with either hCCNT1, hCCNT1.C261Y, or a vector control by transfection. A two-tailed Student’s *t*-test with Welch’s correction was used to compare CCNT1.C261Y cells expressing exogenous hCCNT1 to those expressing hCCNT1.C261Y.

To address Tat activity independently of infection, we next employed a transient reporter assay wherein a firefly luciferase gene under transcriptional control of the HIV-1 LTR (LTR-Luc) was expressed in the presence or absence of Tat in WT and both C261Y cell lines (Fig 3H) (55). When Tat was expressed in the presence of the LTR-Luc construct, we observed a luciferase signal >30-fold higher in the parental cell line relative to either C261Y clone (Fig 3H, compare lane 10 to 11 & 12) confirming severe but not absolute disruption to Tat-driven gene expression, and again significantly rescued when co-expressed with WT human CCNT1 (Fig 3H, compare lane 14 to 17; compare lane 15 to 18; additionally, compare lanes 4-6 to lanes 13-15). Together, these results demonstrated that otherwise permissive human T cells can be rendered resistant to Tat function engendered by a single amino acid substitution at position 261.

### The CCNT1.C261Y modification potently inhibits diverse HIV-1 strains as well as HIV-2 and SIV_agm_

Having confirmed a Tat-specific block using viral reporter assays, we next examined the effect of C261Y on *bona fide* HIV-1 in cultures, under conditions of continuous viral replication and spread. We first infected each cell line with the CXCR4 (X4) tropic HIV-1.NL4-3 or HIV-1.IIIB strains (multiplicity of infection, MOI 0.5) and maintained the cultures for 10 days to allow for multiple rounds of replication. After 10 days, compared to parental cells there was no evidence of infectious virion production in infected CCNT1.C261Y cultures, and we detected a striking, commensurate lack of viral RNA expression (Fig 4A, compare panels ii. to v. & viii. and compare panels iii. to vi. & ix.; quantitation in Fig 4B). To test for potential viral escape, we performed infections with a higher viral dose (HIV-1.NL4-3, MOI ∼10) and maintained the cultures for 30 days to allow time for viral adaptation and outgrowth. No signs of virus amplification following the inoculation were evident, confirming a strong and persistent block (Fig 4C). Because most transmitted strains of HIV-1 are R5-tropic, we also tested transmitted / founder strains HIV-1.CH106.c/2633, HIV-1.RHPA.c/2635, and HIV-1.SUMA.c/2821 in CCNT1 WT and C261Y lines engineered to stably express CCR5 (Fig 4D) (56). Again, compared to the R5-positive parental cells, we observed no evidence of virus replication in R5-positive CCNT1.C261Y cells for 20 days post-infection.

**Fig 4.**
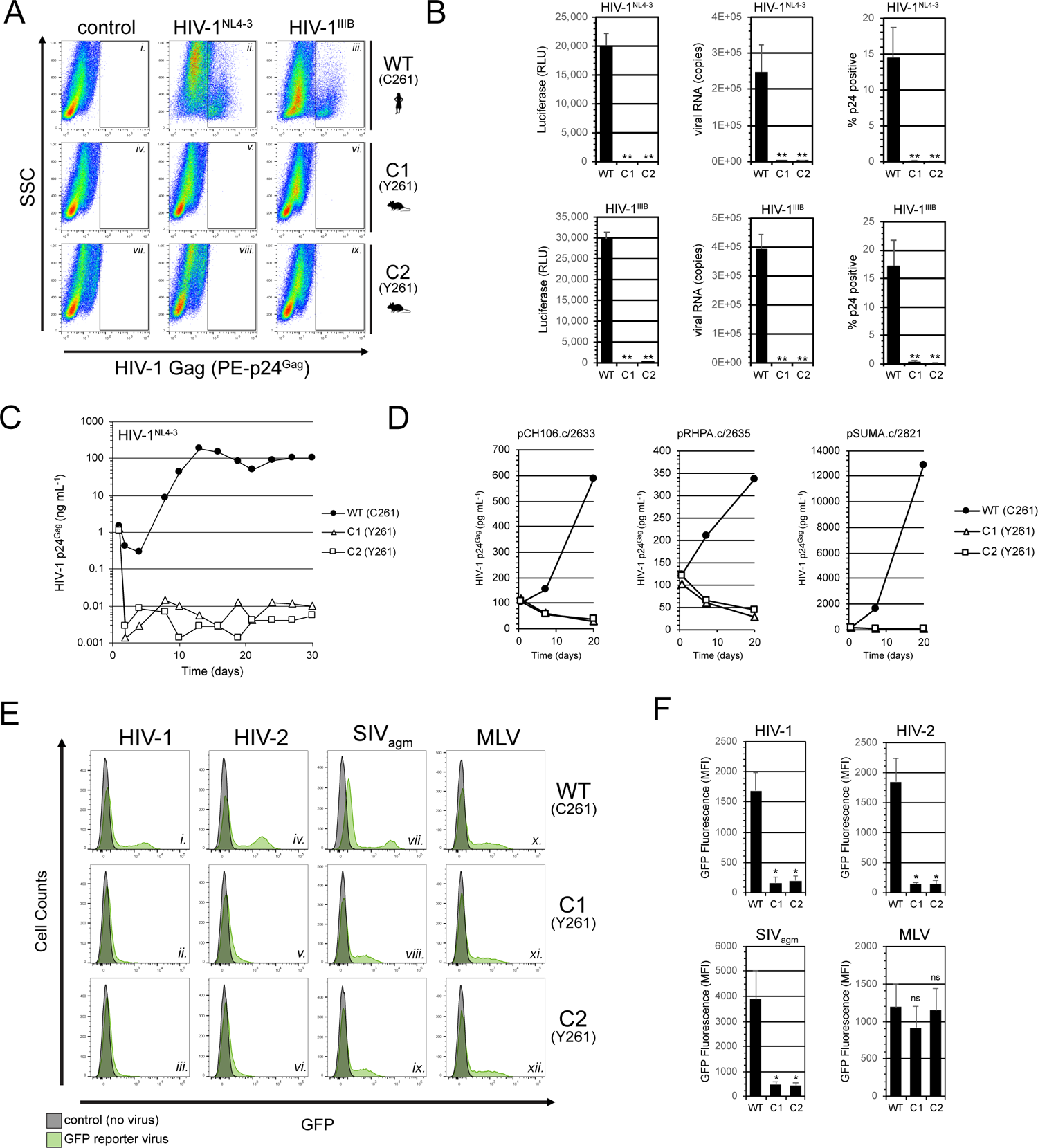
The CCNT1.C261Y modification potently blocks the replication of diverse HIV-1 strains and also inhibits HIV-2 and SIV_agm_ gene expression. (A) CCNT1.C261Y cells are resistant to HIV-1 replication. Representative p24^Gag^ antigen staining of HIV-1 infected cells 10 d post-infection. (B) Quantitative analysis of HIV-1.NL4-3- and HIV-1.IIIB-infected cell cultures 10 d post-infection by supernatant infectivity (TZM-bl luciferase assays, left bar graphs) and viral RNA content (qRT-PCR, middle bar graphs), as well as infected cell populations (p24^Gag^ staining, right bar graphs). A two-tailed Student’s *t*-test with Welch’s correction was used to compare C1 or C2 measurements to wild-type (\*\**p* ≤ 0.01). (C) Cell cultures inoculated with HIV-1.NL4-3 were sampled periodically over 1 month span for p24^Gag^ antigen measurements. (D) Cell cultures inoculated with the CCR5-tropic, transmitted or founder (T/F) clones, HIV-1.CH106, HIV-1.RHPA, or HIV-1.SUMA were sampled periodically over 3-week span for p24^Gag^ antigen measurements. (E) CCNT1.C261Y cells limit viral gene expression for HIV-1 (panels *i.-iii.*), HIV-2 (panels *iv.-vi.*), and SIV_agm_ (panels *vii.-ix.*) but not MLV (panels *x.-xii.*). Flow cytometry histograms from one representative experiment are shown. (F) Mean fluorescence intensities (MFIs) from transduced, GFP-positive cell populations were measured in three independent experiments. A two-tailed Student’s *t*-test with Welch’s correction was used to compare C1 or C2 measurements to wild-type (\**p* ≤ 0.05; ns, not significant).

Because primate lentiviruses code for divergent Tat proteins that similarly drive transcription elongation (57), we tested if CCNT1.C261Y cells would also inhibit transcription using validated GFP-encoded viral vector systems derived from HIV-1, HIV-2, simian immunodeficiency virus from the African green monkey (SIV_agm_) (58,59), and the Tat-independent murine leukemia virus (MLV) as a control. For each vector except for MLV, GFP fluorescence was significantly diminished in the C261Y cell lines compared to the WT parental cells, consistent with the notion that most if not all primate lentiviruses rely on C261 to achieve efficient transcription (Fig 4E and Fig 4F). Taken together, the C261Y substitution appears to be sufficient to achieve remarkably broad-spectrum resistance to primate lentiviruses.

### Infected CCNT1.C261Y cells establish an intrinsic “block-and-lock” provirus inactivation scenario

Given the canonical role of CCNT1 residue C261 in HIV-1 infection, we hypothesized the CCNT1.C261Y cells would be susceptible to HIV-1 infection but yield forced latency due to Tat inactivation. To test this hypothesis, we infected WT and C261Y cell lines with an established latency-tracking HIV-1 reporter virus, R7GeMC, that encodes two reporter cassettes engineered into the *nef*-coding position; an *egfp* gene responsive to the viral LTR promoter and Tat, and a *mcherry* gene under control of a constitutively-active EF1α promoter (Fig 5A; assay depiction in Fig 5B) (60). At 24 hours post-infection, we observed equivalent numbers of mCherry-positive (∼15%) cells for both the CCNT1.C261Y and parental control cells, confirming that HIV-1 entry is not altered in CCNT1.C261Y cell lines and that the antiviral effects occur post-integration (Fig 5C).

**Fig 5.**
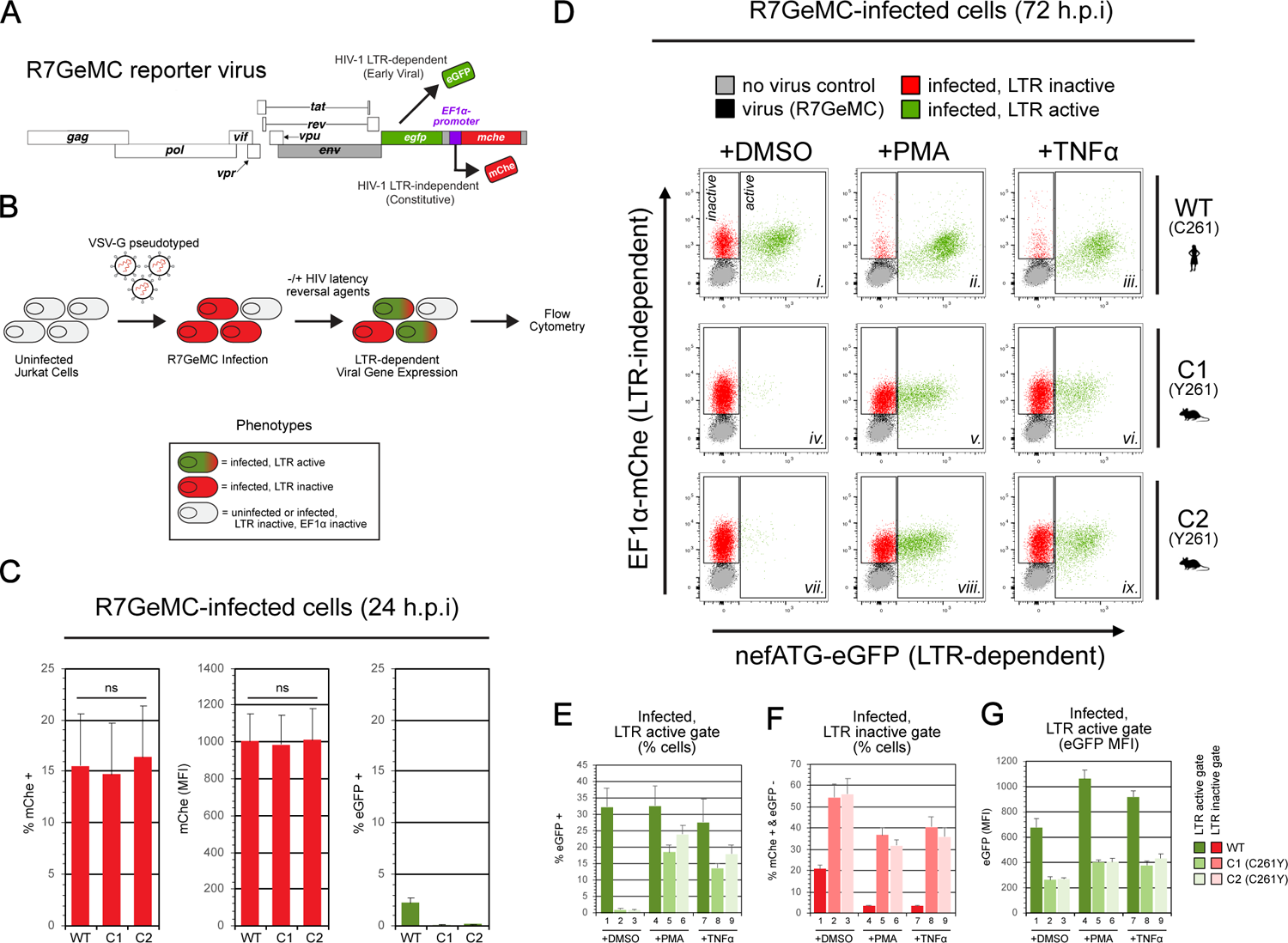
CCNT1.C261Y can be infected by HIV-1 but effectively establish an intrinsic “block-and-lock” provirus inactivation scenario. (A) Virus genome diagram of R7GeMC, an established two-color reporter virus (60) designed to discern transcriptionally-active or inactive HIV-1 proviruses. LTR-dependent viral gene expression is reported by eGFP. Inactive proviruses are reported by mCherry, encoded downstream of the constitutively-active EF1α promoter. Grey rectangles denote inactivated *env* and *nef* genes to limit virus replication to a single cycle. (B) R7GeMC reporter assay schematic. Each cell line was inoculated with VSV-G pseudotyped R7GeMC. 24 h post-inoculation, cells were sampled for fluorescence measurements and the remaining were washed to remove input virus. Cells were split into equivalent cultures and treated with HIV-1 latency reversal agents. 72 h post-inoculation, cells were harvested for fluorescence measurements. (C) Wild-type and CCNT1.C261Y cells are similarly susceptible to R7GeMC. 24 h post-inoculation, eGFP- and mCherry-positive cells and mean fluorescence intensities (MFIs) were measured. A two-tailed Student’s t-test with Welch’s correction was used to compare measurements for C1 or C2 to wild-type (ns, not significant). (D) CCNT1.C261Y cells inhibit integrated R7GeMC proviruses from escaping viral latency, even in the presence of LRAs. Overlaid dot plots of the four indicated cell populations harvested 72 h post-inoculation. (E) Percentage of eGFP-positive infected cells, indicating cells exhibiting active, LTR-dependent viral gene expression (Fig 5D, “active” gate percentages). (F) Percentage of infected mCherry-positive, but transcriptionally inactive eGFP-negative cells, indicating cells with integrated proviruses that lack LTR-dependent viral gene expression (Fig 5D, “inactive” gate percentages). (G) eGFP mean fluorescence intensity (MFI) in the transcriptionally-active infected cell populations (Fig 5D, eGFP MFI in “active” gates).

A recently described HIV-1 functional cure strategy termed “block-and-lock”, aims to use drugs to silence HIV-1 Tat-driven gene expression (“block”) long enough for cell epigenetic modifications to progressively restrict the capacity of HIV-1 to reactivate (“lock”) should drug therapy be discontinued (61). To test the hypothesis that CCNT1.C261Y cells effectively represent a cell-intrinsic “block-and-lock” scenario, we tested the responsiveness of R7GeMC-infected WT and C261Y cells to latency reversal agents (LRAs) that act by potentiating basal, Tat-independent viral gene expression. One day following R7GeMC infection, cultures were washed to remove input virus and treated with the LRAs phorbol 12-myristate 13-acetate (PMA; 10 ng mL^-1^), tumor necrosis factor alpha (TNFα; 10 ng mL^-1^), or with DMSO as a vehicle control. In the absence of LRAs, wild-type cells harbored >28-fold more eGFP-positive cells relative to either CCNT1.C261Y cell line, demonstrating that CCNT1.C261Y cells harbored a significantly greater proportion of inactive proviruses (Fig 5D, compare “active” gate in panel i. to iv. & vii; quantification in Fig 5E; conversely, compare “inactive” gate in panel i. to iv. & vii; quantification in Fig 5F). PMA and TNFα treatment increased the proportion of mCherry- and eGFP-positive CCNT1.C261Y cells (compare “active” gate in panel iv. to v. & vi; and vii. to viii. & ix.; quantification in Fig 5E), however, the net viral gene expression was highly attenuated relative to untreated wild-type cells (compare “active” gate in panel ii. to v. & viii; and iii. to vi. & ix.; quantification in Fig 5E, Fig 5F, and Fig 5G). In sum, the CCNT1.C261Y modification enforces a state of viral latency in human T cells by silencing HIV-1 Tat and, even at early time points post-infection, and severely restricts the potential for viral reactivation in the presence of LRAs, directly achieving the defining characteristics of a “block-and-lock” scenario.

### Editing CCNT1.C261Y causes no discernible effects on p-TEFb activity or major effects on transcription in HIV-negative cells

Finally, the capacity of CCNT1.C261Y cells to achieve cell-intrinsic HIV-1 restriction in the absence of apparent effects on cell viability prompted us to investigate the potential for impacts of the CCNT1.C261Y substitution on p-TEFb complex formation or activity in the context of the cellular host (and independent of HIV-1 infection). The p-TEFb complex is conserved in eukaryotes and regulates transcription elongation for a substantial proportion of RNA polymerase II dependent genes (62–64). While the role for CCNT1 residue 261 in HIV-1 transcription was clear, any effects of targeting C261 on cellular gene expression would need to be considered should this residue/region be considered in the context of an antiviral strategy.

We first examined CCNT1-CDK9 interactions in WT and CCNT1.C261Y cells by co-immunoprecipitation (co-IP) analysis. Comparable amounts of CDK9 were pulled down by CCNT1 or CCNT1.C261Y from respective cell lysates, indicating that p-TEFb complexes were unperturbed by the substitution (Fig 6A, representative blot in Fig 6B). Previous studies had demonstrated that treatment of cells with an RNA polymerase II inhibitor, 5,6-dichlorozo-1-Δ-D-ribofuranosylbenzimidazole (DRB), leads to a shift of p-TEFb from a high molecular weight complex to a low molecular weight complex, reflecting the release of p-TEFb from its transcriptionally-repressive 7SK RNA scaffold (65,66). Upon DRB-treatment, we observed an induction of p-TEFb in CCNT1.C261Y cells (*i.e.*, a decrease in the inactive form of p-TEFb and increase in the active form) comparable to that observed in parental cells (Fig 6C, representative blot in Fig 6D). Thus, p-TEFb responses to signal-induced stimuli appeared to be intact in CCNT1.C261Y cells.

**Fig 6.**
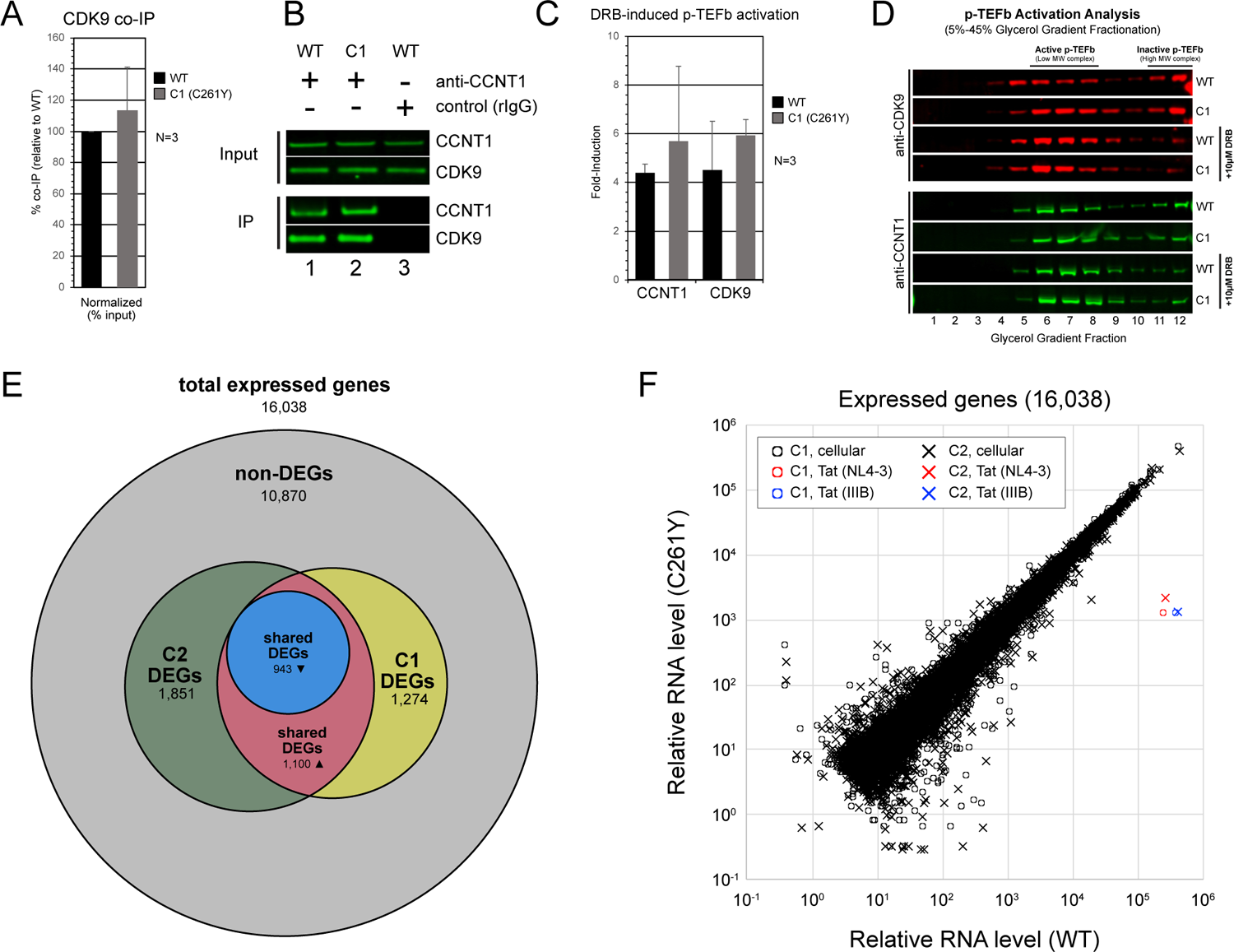
Analysis of p-TEFb and cellular transcriptomes in CCNT1.C261Y cells. (A) CCNT1.C261Y does not alter CDK9-CCNT1 interactions. Protein complexes purified from lysates using CCNT1 antisera were analyzed for CDK9 protein levels. Recombinant IgG (rIgG) was used to control for immunoprecipitation (IP) specificity. CDK9 levels in IP complexes were measured by semi-quantitative western blot. Error bars represent the standard deviation from the mean for three independent experiments. (B) Representative western blot for data presented in Fig 6A. (C) CCNT1.C261Y does not change DRB-induced p-TEFb activation responses. Parental and CCNT1.C261Y clone C1 were cultured in the presence or absence of 10 µM DRB and lysed. Cellular lysates were subjected to glycerol gradient fractionation (5%-45%) and the levels of CCNT1 and CDK9 protein were measured by semi-quantitative western blot. Error bars represent the standard deviation from the mean for three independent experiments. (D) Representative western blot for data presented in Fig 6C. (E) Venn diagram of the summarized differential gene expression analysis comparing CCNT1.C261Y and wild-type transcriptomes (DESeq2). Total expressed genes (16,038) were defined by a baseMean count value of ≥5. Differentially-expressed genes (DEGs, 5,168) were defined by an adjusted p-value (p*) ≤ 0.05. DEGs unique to C1 (1,274), C2 (1,851), or shared in both (2,043) are defined. DEGs that are up-regulated (1,100) or down-regulated (943) in both C1 and C2 compared to wild-type cells are defined. (F) Relative levels of cellular and Tat-dependent RNA in wild-type and C261Y cell lines. Relative RNA abundance (normalized read counts) for all expressed genes (16,038) in uninfected cells are plotted (C1, circles; C2, crosses). Tat-dependent HIV-1 transcript levels (RT-qPCR copy number; from Fig 4B) are embedded for context (red, NL4-3; blue, IIIB).

To employ a more global approach, we next sequenced and compared the transcriptomes of uninfected WT and both C1 and C2 CCNT1.C261Y cell clones, performing differential gene expression analysis using DESeq2 with a low stringency threshold (adjusted p-value, p*, ≤ 0.05, baseMean count value of ≥ 5). This analysis confirmed the majority (>67%) of differentially expressed genes (DEGs) to be unaffected by the C261Y modification (Fig 6E), with most of the remaining DEGs (19.5%) specific to either the C1 or C2 clone. 12.7% of DEGs were observed in both C261Y cell lines relative to WT, indicating genes potentially linked to the C261Y substitution. However, the magnitude of these changes was very low (typically <2-fold) compared to the hyperstimulatory effects of C261 on Tat-dependent gene expression (>100-fold in replication experiments) (Fig 6F), underscoring the relative specificity of CCNT1.C261 regulating viral versus cellular transcription.

Analogous to HIV-1 Tat, the cellular bromodomain-containing protein 4 (BRD4) recruits p-TEFb to target cellular promoters to exert control at a transcription elongation step (Fig 7A) (67), with Tat thought to compete with BRD4 for p-TEFb binding (67,68). Therefore, we surveyed the published literature to compile lists of experimentally defined genes reported to be regulated by either CCNT1 (69–71) or BRD4 (72–79), and asked if these gene sets were affected by the C261Y modification (Fig 7B), using the same low stringency statistical threshold described above. Based on these criteria, only two potential CCNT1-associated genes (DTX4 and PCOLCE2) and two BRD4-associated genes (MCTP2 and IL15RA) differentially expressed more than 2-fold in both CCNT1.C261Y cell clones. Accordingly, the strong majority of genes associated with BRD4 and CCNT1 are unchanged in CCNT1.C261Y cells, supporting the conclusion that, unlike Tat-dependent viral gene activation, cellular gene regulation in Jurkat T cells is not largely dependent on CCNT1 residue 261.

**Fig 7.**
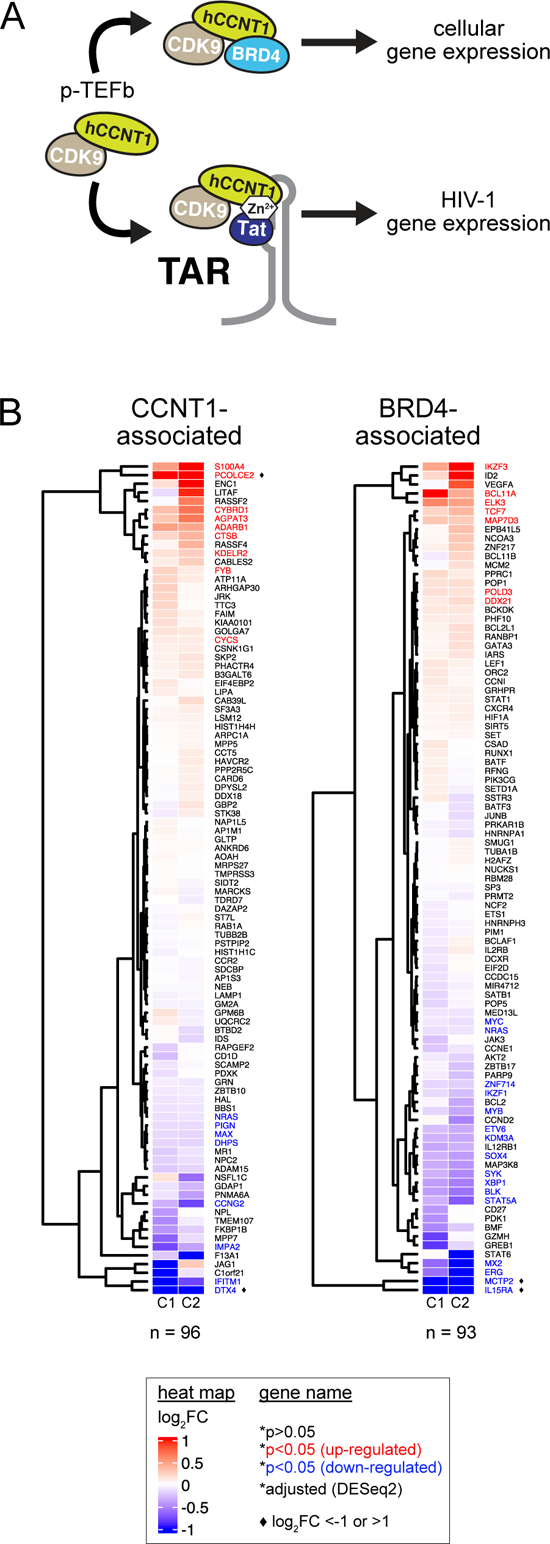
CCNT1- and BRD4-associated genes are largely unperturbed in CCNT1.C261Y cells. (A) Cartoon depiction of p-TEFb-dependent gene expression using either BRD4 or Tat/TAR complexes. (B) CCNT1- and BRD4-associated gene expression is largely preserved in CCNT1.C261Y cells. Heat maps displaying the log_2_FC of CCNT1 (n = 96, left) or BRD4 (n = 93, right) associated genes in the C261Y clones compared to wild-type. Individual genes sharing positive (red) or negative (blue) log_2_FC values were color-coded (adjusted p-value (p*) ≤ 0.05 (DESeq2) in both C1 and C2 cell lines). Diamond symbols (♦) denote differentially expressed genes with log_2_FC of >1 or <-1.

## Discussion

In this study, we engineered human T cells using CRISPR/Cas9 genome editing to test the hypothesis that minor, “rodent-inspired” gene modifications can create a human T cell environment exhibiting intrinsic resistance to HIV-1 early and/or late stage viral gene expression. We provide proof-of-concept for manifesting known blocks to Tat-and Rev-dependent HIV-1 gene expression in human Jurkat T cells using CRISPR/Cas9-based editing of relevant species-specific loci encoded within the endogenous *CCNT1* and *XPO1* genes, respectively. Furthermore, we validated the strength and specificity of the Tat block by isolating two HIV-1-refractory cell clones (C1 and C2) that contained prescribed, homozygous knock-in edits encoding a single non-synonymous mutation in endogenous *CCNT1*, resulting in the C261Y substitution.

Because our study relied heavily on measuring changes to HIV-1 Tat- and/or Rev-dependent gene expression in infected cell populations, we created the E-R-/2FP dual reporter assay to rapidly detect and measure defects to Tat- and Rev-dependent gene expression. In Fig 1, we validated this assay by screening non-human mammalian cell lines predicted to pose genetic blocks to HIV-1 Tat and Rev activity (38,80). As expected, rodent cell lines were largely refractory to HIV-1 Tat and/or Rev function. However, that Rat1 and CHO cells exhibited substantial early, Rev-independent viral gene expression was surprising considering that they encode the non-permissive CCNT1 tyrosine residue at the 261-equivalent position. The competency for early viral gene expression in these cell lines raises the possibility that there are other cell-intrinsic factors that overcome or offset the HIV-refractory effects of the Y261 allele, at least in some cell types. Cat (CRFK) cells are also of interest to us going forward because they supported Rev function despite the cat XPO1 “patch” domain differing from that of the human ortholog at two of the three known relevant species-specific residues.

Recent advances to precision CRISPR/Cas9 genome editing provided us with the means to investigate the potential to introduce species-specific Tat and Rev blocks into human T cells. Unlike HIV-refractory Δ32 *CCR5* alleles, the C261Y and P411T/M412V/F414S substitutions we engineered had not been documented in primates; and thus, at the outset, it was unclear whether human cells would tolerate these changes. Indeed, *CCNT1* and *XPO1* have only experienced modest changes throughout mammalian speciation, with the mouse and human orthologs of CCNT1 and XPO1 exhibiting 89% and 98% amino acid identity, respectively. Considering such strong negative selection, the significance of these HIV-relevant species-specific residues to basic cellular functions (if any) remains unknown. Using our CRISPR/Cas9 knock-in designs, we successfully achieved targeted editing of both *CCNT1* and *XPO1*, observing viral gene repression consistent with disruption of Tat and Rev function in viable cell populations. We also readily isolated seamlessly edited cell clones encoding exclusively CCNT1.C261Y, allowing us to confirm unequivocally that the modification is tolerated in this context. However, attempts to isolate comparably edited homozygous XPO1.P411T/M412V/F414S cells were more challenging and are ongoing.

Using the CCNT1.C261Y cell clones, we confirmed that the C261Y modification imposed a severe block specific to Tat-dependent viral gene expression rescued by exogenous delivery of wild-type CCNT1, and potently inhibited spreading HIV-1 replication (Figs 3 and 4). Given the apparent tolerance and anti-HIV effects of the C261Y substitution in human T cells, we considered this knock-in approach in the context of a “block-and-lock” functional cure scenario (81), employing a latency reporter system (60) to demonstrate that the CCNT1.C261Y cells remain susceptible to HIV-1 infection but are markedly refractory to drug-induced viral reactivation (Fig 5). Akin to this block, a recently-described small molecule, didehydro-cortistatin A (dCA) has shown promise as a viral transcription inhibitor disrupting Tat/TAR interactions (82). Valente and colleagues recently isolated dCA-resistant HIV-1 clones with dCA resistance mapped to mutations in the viral LTR, *nef* or *vpr* genes; all of which were suggested to promote basal, Tat-independent transcription and viruses lost the ability to enter viral latency (83). To date, we have no indication of viral resistance to CCNT1.C261Y in virus replication assays (Fig 4), although these efforts will be ongoing.

In Fig 6, we showed that human T cells bearing CCNT1.C261Y limit the activity of not only HIV-1 Tat, but also the Tat proteins of HIV-2 and SIV. These results emphasize how fundamental the Tat-C261 interaction must be to the replicative success of primate lentiviruses and suggests that any intervention strategy that targets the crucial CCNT1.C261 residue will be broad-spectrum, likely affecting all HIV-1 and HIV-2 strains and subtypes. Moreover, because this mechanism extends to SIV, the block may be testable in non-human primate model systems. Interestingly, a subset of non-primate lentiviruses are thought to be less dependent on CCNT1 C261 (e.g., bovine immunodeficiency virus, BIV) (84,85) so that additional comparative structure-function studies of Tat-CCNT1 interactions seem warranted.

Aside from their crucial roles in HIV-1 Tat and Rev function, little is known regarding the relevance, if any, of CCNT1 residue 261 or XPO1 residues 411, 412, and 414 to cellular processes; a key consideration should these protein domains be targeted in the context of antiviral strategies. CCNT1.C261Y clones exhibited no significant changes to rates of cell proliferation or basal *CCNT1* expression (Fig 5), confirming that the C261Y modification is not profoundly detrimental, at least in cell lines. In Fig 6, we further investigated the C261Y modification with experiments designed to interrogate endogenous p-TEFb complexes and their response to stimuli, and performed a comprehensive transcriptomic analysis of CCNT1.C261Y cell lines. CDK9 co-IP experiments indicated that the steady-state levels and interactions between CCNT1 and CDK9 were unperturbed relative to the parental cells. *CCNT1* knock-out cells and mice depleted of CCNT1 have been reported with notable changes to the expression of *CDK9* and other transcription factors (86,87). However, our glycerol gradient analysis indicated similar expression of CDK9 for both wild-type *CCNT1* and *CCNT1.C261Y* cells, as well as successful release of CDK9 from high molecular weight complexes in both cell backgrounds in response to DRB treatment (66,65). Further, transcriptomic analysis using RNA-seq demonstrated that the majority of CCNT1- and BRD4-dependent genes were unperturbed in the *CCNT1.C261Y* background. Taken together, the effects of *CCNT1.C261Y* are largely virus-specific, being far more severe on Tat-dependent gene expression relative to any subset of cellular genes.

It remains highly interesting to us and unexplained why rodents code for variation in these otherwise conserved *CCNT1* and *XPO1* genes, with these variations prohibiting HIV-1 gene expression in a synergistic fashion. The functional consequences of this constellation of polymorphisms on modern lentiviruses including HIV-1 but also HIV-2 and SIV_agm_ (Fig 4) is remarkable. We propose that harnessing these species-specific polymorphisms to engineer cell-intrinsic resistance barriers into human T cells will provide targeted means to study these selective pressures, and should also be considered in the context of functional-cure based antiviral strategies.

## Materials & Methods

### Cell culture

Human T lymphocyte (Jurkat E6-1), human embryonic kidney (293T), human cervical carcinoma (HeLa), mouse fibroblast (3T3), and Chinese hamster ovary (CHO) cell lines were obtained from the American Type Culture Collection (ATCC). African green monkey osteosarcoma (Cos7), chicken fibroblast (DF-1), cat kidney (CRFK), mouse fibroblast (L cells, and M. dunni) and mouse myoblast (C2C12) cell lines were a kind gift of Dr. Michael Malim (King’s College London). The Rat fibroblast (Rat1) cell line was a kind gift of Dr. Robert Kalejta (University of Wisconsin-Madison). The chimpanzee skin fibroblast (WES) cell line was a kind gift of Dr. Beatrice Hahn (University of Pennsylvania). The TZM-bl reporter cell line was a kind gift of Dr. John Kappes and Dr. Xiaoyun Wu and obtained from the HIV Reagent Program (NIH). CHO, Jurkat and Jurkat-derived cell lines were cultured in Roswell Park Memorial Institute (RPMI) 1640 supplemented to 10% Fetal Bovine Serum (FBS), 1% penicillin-streptomycin and 1% L-glutamine (Sigma). All other established cell lines were cultured in Dulbecco’s Modified Eagle’s Medium (DMEM) supplemented to 10% FBS, 1% L-glutamine, and 1% penicillin-streptomycin. Cell cultures were maintained at 37°C and 5% CO_2_ in a humidified incubator.

### Plasmids

The E-R-/2FP dual reporter virus construct is derived from the E-R-/mCherry construct described elsewhere (88). Using overlapping PCR and standard molecular cloning procedures, cDNA sequences encoding the mVenus fluorescent protein with flanking glycine-rich linker segment at the 5’-(GGSGGTR) and 3’-(TRGGSGG) ends were fused in frame with the HIV-1 *gag* sequence between matrix and capsid coding regions (89). E-R-Tat-/2FP was generated by overlapping PCR using the primers 5’-CCT AGA CTA GAG CCC TGG AAG CAT CCA TGA AGT TAA CCT AAA ACT G-3’ and 5’-GGC TCT AGT CTA GGA TCT ACT GGC TCC GTT TCT TGC TC-3’. E-R-Rev-/2FP was generated by subcloning using E-R-Rev-/YFP described elsewhere (90). The lentiviral packaging plasmid psPax2 was a kind gift from Dr. Didier Trono (plasmid #12260, Addgene). The pVSV-G expression construct is described elsewhere (91). Retroviral vector plasmids encoding *CCNT1* variants were generated by subcloning modified *CCNT1* cDNA sequences fused to cDNA encoding the FLAG-epitope tag (DYKDDDDK) into the pELE88 vector (80) using BglII and NotI restriction enzyme sites. Retroviral vector plasmids encoding GFP or CCR5 were generated by subcloning cDNAs into MIGR1-derived vectors (92) using vector-encoded BglII and XhoI restriction enzyme sites. The retroviral packaging plasmid pMD.GagPol was a kind gift from Dr. Richard Mulligan. For the Tat/LTR reporter assay, the construction of the *gaussia* luciferase (gLuc) HIV-1 LTR reporter construct is described elsewhere (55). The pEV280 Tat-FLAG expression construct was a kind gift of Dr. Melanie Ott. Additional plasmids used for experimental controls include pmCherry-C1 (Clontech Laboratories Inc., Mountain View, CA, USA), cpTK-Cluc (New England Biolabs, Ipswich, MA, USA), pBluescript (Stratagene, La Jolla, CA, USA), and pSEAP (Clontech Laboratories Inc.). Plasmids encoding the infectious molecular clones of HIV-1, pCH106.c/2633, pRHPA.c/2635, and pSUMA.c/2821 were kind gifts of Dr. John Kappes and Dr. Christina Ochsenbauer and obtained from the HIV Reagent Program (NIH). The R7GEmC reporter virus (60) was a kind gift of Dr. Vincenzo Calvanese and Dr. Eric Verdin and obtained from the HIV Reagent Program (NIH). The GFP-encoding and *env*-deficient HIV-1.NL4-3, HIV-2.ROD, and SIV_agm_.Tan-1 reporter virus clones are described elsewhere (58,59) and were a kind gift of Dr. Paul Bieniasz.

### Virus preparation

To prepare virus stocks, 1 x 10^6^ 293T producer cells were plated in 10 cm plasmid tissue culture treated dishes. 24 h post-plating, cells were transfected using polyethylenimine (PEI; catalog no. 23966. Polysciences, Inc., Warrington, PA, USA) followed by culture medium exchange 4 h post-transfection. 48 h post-transfection, cell culture supernatants were 0.45um filtered, aliquoted, and stored at −80°C. For E-R-/2FP reporter virus stocks, plasmid mixes contained 4 μg E-R-/2FP variant, 4 μg psPax2, and 1 μg pVSV-G. For R7GeMC reporter virus stocks, plasmid mixes contained 9 μg R7GeMC and 1 μg pVSV-G. For GFP-encoding primate lentivirus reporter stocks, plasmid mixes contained 9 μg each E-GFP vector (HIV-1.NL4-3, HIV-2.ROD, or SIV_agm_.Tan) and 1 μg pVSV-G. For MLV-based retroviral vector stocks encoding GFP, CCNT1, or CCNT1.C261Y, plasmid mixes contained 4 μg each retroviral vector plasmid, 4 μg pMD.GagPol and 1 μg pVSV-G. For replication-competent HIV-1 virus stocks, 10 μg plasmid encoding each infectious HIV-1 molecular clone were used. When necessary, reporter virus stocks were titrated on Jurkat E6-1 cells to establish normalized infectious doses. Replication-competent HIV-1 stocks were normalized by p24^Gag^ levels determined by ELISA (PerkinElmer Life Sciences, Inc., Boston, MA, USA).

### E-R-/2FP infection assays

For infections with suspension cell lines, 1 x 10^6^ cells were plated in each well of a 6-well dish and infected immediately with equivalent doses of E-R-/2FP virus stock. For infections with adherent cell lines, the indicated cell line was plated to 30-50% confluency in each well of a 6-well dish and infected 24 h post-plating as above. Nevirapine stock was obtained from the HIV Reagent Program (NIH) and added immediately prior to infection to a final concentration of 1 μM where indicated. 48 h post-infection, cells were harvested, washed in PBS, fixed with 4% reconstituted paraformaldehyde (PFA), and analyzed by flow cytometry. Flow cytometry was performed using either a LSRII (BD Biosciences, Mississauga, ON, Canada) or an Attune NxT (Thermo Fisher Scientific, Waltham, MA, USA). Flow cytometry data analysis was performed using FlowJo software (FlowJo, LLC, Ashland, OR, USA). For CCNT1 rescue experiments, 5 x 10^5^ cells were transduced by spinoculation (93) with retroviral vectors encoding CCNT1, CCNT1.C261Y, or vector only control in the presence of 5 μg/mL polybrene (Sigma) prior to plating. Following transduction, cell cultures were diluted 1:1 with fresh culture medium volume, plated in 6-well tissue culture treated dishes and cultured for 3 d. Cells were then infected, harvested, and analyzed as described above.

### Genome engineering of Jurkat E6-1 cells

Alt-R^TM^ recombinant S.p. Cas9 nuclease-3NLS (IDT, Integrated DNA Technologies, Coralville, IA, #1074181), custom Alt-R^TM^ CRISPR-Cas9 crRNAs (IDT), ATTO^TM^-550 labeled Alt-R^TM^ tracrRNA (IDT, #1075928), and Alt-R^TM^ Cas9 electroporation enhancer reagents were prepared and ribonucleoprotein (RNP) complex assembly was performed according to the manufacturer’s instructions. *CCNT1* crRNA: 5’-TGC TTC ATT GCA GGC ATG CG-3’. *XPO1* crRNA: 5’-TCT GTT ACC TTG AAT AAC AT-3’. The indicated 119-nt single-stranded oligodeoxynucleotide (ssODN) donor template for homology-directed repair (HDR) was custom prepared (Sigma) and resuspended in TE buffer. *CCNT1* ssODN donor template: 5’-GTG TTT TTT TAT TTA GTA AAT TAC CTA AGT AAA GAG ATG CTA TTT GCT TCA TTG CAG GCG TAC GAA GCT GCC AAG AAA ACA AAA GCA GAT GAC CGA GGA ACA GAT GAA AAG ACT TCA GA-3’. *XPO1* ssODN donor template: 5’-TGC TTT CTG GAA GTC AAC ATT TTG ATG TTC CTC CCA GGA GAC AGC TGT ATT TGA CTG TGT TAT CAA AGG TAA CAG AGC GGT TGG TTG AGT GTT CTT CCT GTT GCA TAC TGT GGT TTT GA-3’. RNP complexes and ssODNs were delivered to Jurkat E6-1 cells using the Neon® Transfection System and Neon® Transfection 10uL Kit (Thermo Fisher Scientific) according to manufacturer’s instructions. Electroporation parameters were 1600V, 10ms pulse width, 3 pulses and cells were cultured post-electroporation in antibiotic-free media (RPMI 1640 supplemented with 10% FBS). 24 h post-electroporation, cells were bulk-sorted by fluorescence-associated cell sorting (FACS) to concentrate ATTO^TM^-550 positive cells in antibiotic-replete media (RPMI 1640 supplemented with 10% FBS and 1% penicillin-streptomycin-L-glutamine). The Jurkat E6-1 cell line was selected for gene knock-in editing because the chromosomal regions encoding *CCNT1* and *XPO1* were reportedly normal as determined by karyotype analyses (ATCC).

### CRISPR knock-in screening

Cells from each CRISPR treatment were pelleted, washed in PBS, and resuspended in 10 uL of 1X Green GoTaq Flexi Buffer (Promega, Madison, WI, USA) and stored at −80°C. Frozen cell samples were thawed and subjected to proteinase K treatment (65°C for 1 h; 95°C for 15 min; 4°C hold) to yield genomic DNA template mixture. During hold step, 40 μL reagent mix containing 1X Green GoTaq Flexi Buffer, 0.5 μM each screening primer, 0.5 μM dNTP mix, 1.5 mM MgCl_2_, and 2.5 U GoTaq DNA polymerase (GoTaq Green Flexi Kit, Promega) was added to 10 μL genomic DNA template mixture yielding 50 μL total reaction volume. Target-dependent screening primers are as follows: *CCNT1* forward screening primer: 5’-TGA GAT TAG AAG TAG GCT TGA GAG G-3’. *CCNT1* reverse screening primer: 5’-GCT AAA TTC TCA CTA GTC CGA TGA C-3’. *XPO1* forward screening primer: 5’-TTC TCT CCT CTG TGA TGG TAC ATT T-3’. *XPO1* reverse screening primer: 5’-TCA AGA TTG TAG TGA GCT ATG ACC A-3’. Optimized PCR cycling conditions yielded 1 kb-sized PCR amplicons for each target. Where indicated, prepared PCR amplicons were submitted for Sanger sequencing (Functional Biosciences, Madison, WI, USA) for analysis. Sequencing chromatograms were prepared using SnapGene software (GSL Biotech, San Diego, CA, USA). Restriction enzyme digests of PCR amplicons were carried out following the manufacturer’s recommended reaction composition with BsiWI or SacI for *CCNT1* or PvuII or XbaI for *XPO1*. Predicted *CCNT1* SacI digestion products: 804 bp and 196 bp. Predicted *CCNT1* BsiWI digestion products: 712 bp and 288 bp. Predicted *XPO1* XbaI digestion products: 679 bp and 298 bp. Predicted *XPO1* PvuII digestion products: 497 bp and 480 bp.

### E-R-/2FP FACS analysis

For infections of heterogeneous populations of CRISPR-treated Jurkat cells, equivalent numbers of unfixed and infected cells from each gate (>10,000 per gate; gate scheme shown in Fig 2C) were analyzed by flow cytometry and sorted by FACS using a FACSAria cell sorter (BD Biosciences) instrument. Bulk-sorted cells were screened for CRISPR knock-in as described above. Corresponding flow cytometry measurements were analyzed as described above.

### SDS-PAGE and Immunoblotting

To measure endogenous CCNT1 protein levels, equivalent numbers of each Jurkat cell line were pelleted, washed with PBS, and in lysed using radioimmunoprecipitation assay (RIPA) buffer (10 mM Tris-HCl [pH 7.5], 150 mM NaCl, 1 mM EDTA, 0.1% SDS, 1% Triton X-100, 1% sodium deoxycholate) containing complete protease inhibitor cocktail (Roche). Cell lysates were prepared for immunoblot by sonication and clarified supernatants were boiled with 2X dissociation buffer (62.5 mM Tris-HCl [pH 6.8], 10% glycerol, 2% sodium dodecyl sulfate [SDS], 10% ϕ3-mercaptoethanol) at a 1:1 ratio prior to polyacrylamide gel electrophoresis (PAGE) and immunoblot. Prepared cell lysates were subjected to SDS-PAGE using 10% polyacrylamide gels and a conventional Tris-glycine buffering system. Following electrophoresis, resolved lysates were transferred to 0.2μm pore size nitrocellulose membranes (GE Healthcare) for immunoblot. Nitrocellulose membranes were blocked in blocking solution (PBS containing 0.1% Tween-20 (PBS-T) and reconstituted nonfat dry milk at a 2% final concentration). Following incubation in the blocking solution, membranes were incubated in primary antibody solutions consisting of fresh blocking solution containing CCNT1 antisera (Santa Cruz Biosciences, Dallas, TX, USA, sc-398695; Cell Signaling Technologies, Danvers, MA, USA D1B6G), CDK9 antisera (Bethyl Laboratories, Montgomery, TX, USA, A303-493A), or HSP90 antisera (Santa Cruz Biosciences, sc-7947; sc-13119). Following incubation in primary antibody solution, membranes were washed three times with PBS-T and incubated in secondary antibody solutions consisting of fresh blocking solution containing either anti-mouse or anti-rabbit secondary antisera conjugated to either IRDye680 or IRDye800 (LI-COR Biosciences, Lincoln, NE, USA) infrared fluorophores. Following incubation in secondary antibody solution, membranes were washed three times in PBS-T and analyzed by quantitative immunoblot using an Odyssey Fc instrument (LI-COR Biosciences).

### Cell Proliferation Assay

5 x 10^4^ cells from each Jurkat line were plated in each well of a 12-well tissue culture treated dish containing 1 mL of cell culture medium. At 4 and 6 d post-plating, cultures for each cell line were resuspended, diluted, and mixed at 1:1 concentration with a 0.4% Trypan blue (Sigma) solution. Cells lacking Trypan blue uptake were counted manually using a hemocytometer.

### Transient HIV-1 LTR reporter assays

For each Jurkat cell line, 5 x 10^5^ cells were transfected using the Neon® electroporation system using the electroporation parameters described above per the manufacturer’s instructions. Plasmid mixtures contained 75 ng of HIV-1 LTR reporter constructs *gaussia* luciferase (gLuc) and nanoluciferase, (nLuc, Promega), 250 ng pmCherry-C1 plasmid, 250 ng pTK-cLuc plasmid, 1200 ng CCNT1 or control plasmid (pBluescript or pSEAP), 25 ng pEV280 Tat expression plasmid or control carrier DNA. Total DNA concentrations were maintained at 2,500 ng by vector control plasmids or calf thymus DNA. 24 h post-transfection, 10 μL culture media was removed, diluted with 40 μL PBS and assayed for secreted *gaussia* Luciferase (gLuc) activity by injecting 30 μL coelenterazine solution (Promega) followed by 1.6 sec incubation period and a luminescence integration read time of 1.0 sec. Secreted *cypridina* Luciferase (cLuc) activity from the internal control plasmid was similarly measured using the *cypridina* Luciferase kit (New England Biolabs) according to the manufacturer’s instructions. Fold activation was calculated as the ratio of gLuc: cLuc signals in each well compared to the average of the pBluescript and pSEAP control transfections.

### HIV-1 infection assays

2.5 x 10^6^ cells from each Jurkat cell line were plated in each well of a 48-well plate and infected with HIV variants, NL4-3 or IIIB, at a multiplicity of infection of 0.5. The following day, the inoculum was removed, cells were washed twice with PBS then replenished with fresh culture medium and maintained at normal culture conditions. At 4- and 10-days post infection cells and supernatant were collected for viral RNA quantification, viral outgrowth assays, and flow cytometry staining analysis.

### Viral RNA quantification

Viral RNA measurements were performed using the method described previously (94). From 100 μL of supernatant collected at each time point, viral RNA was extracted using the QIAamp viral RNA mini kit (Qiagen, Hilden, Germany) following the manufacturer’s protocol. SYBR green real-time PCR assay was carried out in a 20μL PCR mixture volume consisting of 10 μL of 2X Quantitect SYBR green RT-PCR Master Mix (Qiagen) containing HotStarTaq DNA polymerase, 0.5 μL of 500 nM each oligonucleotide primer, 0.2 μL of 100X QuantiTect RT Mix (containing Omniscript and Sensiscript RTs), and 8 μL of RNA extracted from samples or standard HIV-1 RNA (from 5 x 10^5^ to 5 copies per 1 ml). Highly conserved sequences on the gag region of HIV-1 were chosen, and specific HIV-1 gag primers were selected. The sequences of HIV-1 gag primers are 5’-CAA TGG CAG CAA TTT CAC CA-3’ and 5’-GAA TGC CAA ATT CCT GCT TGA-3’. Amplification was done in an Applied Biosystems 7500 real-time PCR system (Applied Biosystems, Waltham, MA, USA), and it involved activation at 45°C for 15 min and 95°C for 15 min followed by 40 amplification cycles of 95°C for 15 s, 60°C for 15 s, and 72°C for 30 s. For the detection and quantification of viral RNA, the real-time PCR of each sample was compared with threshold cycle value of a standard curve.

### Viral outgrowth assay in TZM-bl cells

24 h prior to infection, 20,000 TZM-bl cells were plated in each well of a 48-well tissue culture treated plate. The cells were incubated with 10 μL supernatant collected from infected Jurkat cell cultures for overnight incubation at 37°C. The following day, infected cells were washed and fresh culture media was added. 48 h post-infection, luciferase activity in cell lysates was measured using a luciferase assay kit, following the manufacturer’s protocol (Promega).

### p24^Gag^ staining

HIV-infected cells for each Jurkat cell lines were permeabilized using the Cytofix/Cytoperm Fixation/Permeabilization Kit (BD Biosciences) and stained intracellularly using PE-conjugated mouse anti-p24 mAb (clone KC57; Beckman Coulter, Brea, CA, USA, 1:100 dilution). Flow cytometry measurements were performed using a FACSCalibur (BD Biosciences) and data analysis was performed as above.

### HIV-1 replication curves

5 x 10^6^ cells from each Jurkat cell line were infected with 10 ng p24^Gag^-equivalent volumes for the indicated HIV-1 variants in 6 mL antibiotic-replete RPMI. 24 h post-infection, cultures were pelleted, washed three times with 1X PBS, and resuspended in fresh culture medium. Each 6 mL culture was maintained under normal culture conditions and split by removal of 4mL culture (media containing cells) and refreshed with an equal volume of fresh media every 3 days. Where indicated and prior to infection, cells were transduced with a *CCR5*-encoding retroviral vector to confer susceptibility R5-tropic HIV-1 variants. Harvested cultures were processed by 0.45μm-filtration to remove cells and debris and two filtered ∼1mL aliquots were stored at −80°C. As previously described, p24^Gag^ levels were quantified by ELISA periodically (every 1-2 weeks) to monitor supernatant virus particle levels over time.

### R7GeMC infection assays

1 x 10^6^ cells were plated in each well of a 6-well dish and infected immediately with equivalent doses of R7GeMC virus stock. 24 h post-infection, cells were pelleted and washed thrice in PBS, then resuspended in culture medium containing phorbol 12-myristate 13-acetate (PMA, Sigma) or Tumor necrosis factor alpha (TNFα, EMD Millipore, 80054-834) at 10ng mL^-1^ or vehicle control (DMSO) where indicated. 2 d post-treatment, cells were harvested, processed, and analyzed by flow cytometry as described above.

### Retroviral GFP reporter assays

1 x 10^6^ cells were plated in each well of a 6-well dish and infected immediately with equivalent doses of the indicated GFP-encoding reporter virus stock. 24 h post-infection, cells were harvested, processed, and analyzed by flow cytometry as described above.

### Glycerol Gradients

5 x 10^7^ cells for each Jurkat cell line were incubated in the presence of 10 μM 5,6-Dichlorobenzimidazole 1-β-D-ribofuranoside (DRB, Sigma) or DMSO vehicle control for 2 h under normal culture conditions. Cells were then pelleted, washed with chilled PBS, and lysed in 0.5 mL Glycerol Gradient Lysis Buffer (GGLB; 150 mM NaCl, 10 mM KCl, 10 mM MgCl_2_, 10 mM HEPES [pH 7.5], 2 mM Δ-mercaptoethanol, 0.5% NP-40, 1X ProteoBlock protease inhibitor cocktail (Fermentas, Hanover, MD)) for 10 min on ice. Cell lysates were centrifuged to pellet nuclei (20,000 x G for 5 minutes at 4°C) and supernatants were layered atop a 5%-45% glycerol gradient prepared with GGLB. Samples were centrifuged (190,000 x G for 16 h at 4°C) using a TLS-55 rotor. Following centrifugation, fractions were pipetted into fresh tubes, prepared for SDS-PAGE and immunoblotting and analyzed as described above. Fractions 6-8 (active) and 11-12 (inactive) were used to determine DRB responses by calculating fold-induction over untreated controls.

### Co-immunoprecipitation

5 x 10^6^ Tat-expressing cells for each Jurkat cell line were lysed in 500 μL ice cold Tris buffered saline NP-40 (TBSN) [150 mM NaCl, 50 mM TRIS (pH 7.5), 1 mM EDTA, 2 mM Δ-mercaptoethanol, 0.5% NP40, 1X HALT protease inhibitor cocktail (Invitrogen, Carlsbad, CA)] buffer. Cells were lysed on ice for 10 min, nuclei were pelleted (20,000 x G for 10 min at 4°C), and supernatant was transferred to a new tube. 50 μL (10%) of the input material was removed for SDS-PAGE. The remaining sample was immunoprecipitated with anti-CCNT1 antibody (Bethyl Laboratories, A303-499A) or non-specific normal rabbit IgG (Cell Signaling Technologies, 2729S) in the presence of 2,000 gel units micrococcal nuclease (New England Biolabs) for 1 hour. 50 μL magnetic protein A/G beads (Pierce, 88803) were added and the samples incubated for an additional hour. The samples were washed thrice in TBSN and the beads were resuspended in 50 μL SDS-PAGE loading buffer. The samples were boiled, equivalent volumes of Input and IP elution were separated by SDS-PAGE and immunoblotted with mouse anti-CDK9 (Santa Cruz Biosciences, sc-13130) and mouse anti-CCNT1 (Santa Cruz Biosciences, sc-271348) as described above.

### Transcriptome analysis

For each Jurkat cell line, 1.5 x 10^6^ cells in 10mL fresh culture medium were plated in T75 tissue culture treated flasks and maintained under normal culture conditions. 48 h post-plating, cells were pelleted from suspension (400xG for 5 minutes at 4°C) and delicately resuspended in 1mL chilled PBS and transferred to 1.5mL microcentrifuge tube. Cell suspensions were pelleted (400xG, 5 minutes 4°C) and clarified PBS was aspirated. Cell pellets were stored at −80°C. Total RNA was extracted from cell pellets of three biological replicates (*i.e.*, time-separated cultures) using RNeasy Plus Mini Kit (Qiagen) with genomic DNA eliminator columns and resuspended in nuclease-free water according to the manufacturer’s instructions. RNA extracts were stored at −80°C prior to submission to the University of Wisconsin-Madison Biotechnology Center (UWBC) for quality control assessment, library preparation, and sequencing. RNA sample concentrations and quality assessments were measured using a NanoDrop One spectrophotometer (software version 1.4.0) and an Agilent 2100 Bioanalyzer (G2939A; firmware: C.01.069; Assay: Eukaryote Total RNA Nano). Poly(A)-enriched mRNA sequencing libraries were prepared using an Illumina TruSeq® stranded mRNA Library Prep Kit (Illumina, San Diego, CA, USA) according to the manufacturer’s instructions. Samples were loaded onto a MiSeq NanoCell (Illumina) and paired-end RNA sequencing was performed on a NovaSeq 6000 sequencing platform (Illumina). Sequencing was carried out with a read depth target of approximately 30 million reads per sample. Sequencing reads were mapped to human genome assembly hg38 using HISAT2 (95) (version 2.0.5). Annotated feature count tables were prepared using the *htseq-count* tool within HTSeq (96) (version 0.6.0). Differential gene expression (DGE) analyses were performed using DESeq2 (version 1.26.0) (97). Computational resources for RNA sequencing analysis were kindly provided by the Bioinformatics Resource Center (BRC) at the University of Wisconsin–Madison. RNA-seq data have been deposited to the Gene Expression Omnibus (https://www.ncbi.nlm.nih.gov/geo) and can be accessed via GSE209745.

## Acknowledgements

We thank the University of Wisconsin Carbone Cancer Center (UWCCC) Flow Cytometry Shared Resource Facility for training and assistance with flow cytometry, cell sorting, and data analysis. We also thank the Gene Expression Center and DNA sequencing facility at the University of Wisconsin Biotechnology Center (UWBC) for providing library preparation, sample quality control, and next generation sequencing services. We also thank the Malim (King’s College London), Bieniasz (Rockefeller University), Verdin (University of California – Los Angeles), Kappes (University of Alabama), Hahn (University of Pennsylvania), and Kalejta (University of Wisconsin-Madison) laboratories for valuable contributions.

## Supporting Information

**S1 Fig.**
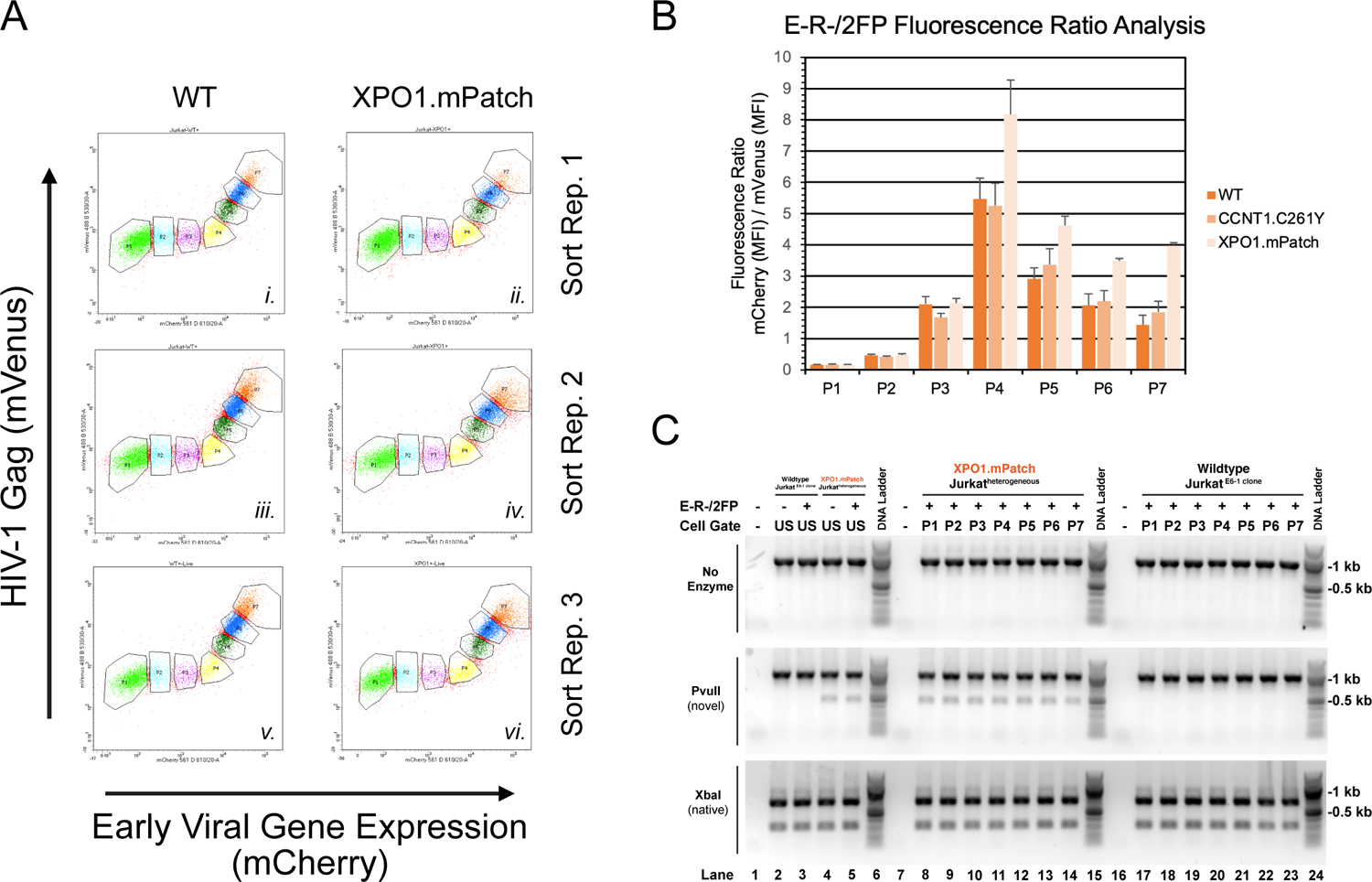
Late viral gene expression in *XPO1*-edited cells requires elevated early viral gene expression levels. (Related to, **Fig 2**. CRISPR/Cas9 knock-in strategy to abrogate Tat or Rev function in human CD4+ T cell lines). (A) Fluorescence profiles of *XPO1*-edited (right column) or control cells (left column) infected with E-R-/2FP. P1-P7 gates, described in Fig 2, are also shown. Each row represents an individual E-R-/2FP infection replicate. (B) Ratio of mCherry to mVenus fluorescence (MFI) in cells inoculated with E-R-/2FP for gates P1-P7 for the *CCNT1*-, *XPO1*-, and control-edited cell populations. Error bars represent the standard deviation from the mean for three independent experiments. (C) Sensitivity of PCR amplicons from E-R-/2FP-inoculated *XPO1*-edited or control cells before and after FACS. PCR amplicons (top row) were digested with PvuII (middle row) or XbaI (bottom row) to screen for amplicons containing gene knock-in for each sorted population (P1-P7, lanes 8-14 and 17-23). Unsorted E-R-/2FP-inoculated or no infection control cell populations (US, Lanes 1-5) were analyzed to demonstrate input amplicon sensitivities to the indicated restriction enzymes.

